# GLproxScape reconstructs spatial chromatin occupancy landscapes from tiled genomic locus proteomics

**DOI:** 10.64898/2026.06.29.735243

**Authors:** Selahattin Can Ozcan, Baris Sergi, Busra Yildirim, Umut Cagiral, Mehmet Gönen, Ceyda Acilan

**Author notes:** These authors contributed equally to this work (Baris Sergi, Busra Yildirim, Umut Cagiral).

## Abstract

Genomic locus proteomics combines proximity labeling with mass spectrometry to identify the proteins associated with user-defined genomic loci. However, per-region enrichment values from tiled guide designs are typically pooled before hit calling, collapsing the latent spatial structure encoded by overlapping measurements. Here, we describe GLproxScape, an R package that treats per-region enrichments as indirect spatial measurements and reconstructs latent chromatin occupancy landscapes through a Gaussian labeling-kernel forward model. Sequence-specific transcription factors are resolved by motif-anchored non-negative least-squares deconvolution against JASPAR or HOCOMOCO position weight matrices, while chromatin regulators which lack defined DNA-binding motifs are inferred as broad occupancy zones, enabling recovery of overlapping members of multi-subunit complexes. Applied to published genomic locus proteomics datasets at the human TERT, MYC, FOXP2, and FOXQ1 loci and the mouse Ripk3 locus, GLproxScape recovered known regulators with predicted positions independently supported by ChIP-Atlas peaks, reconstructed candidate co-binding relationships, and identified chromatin complexes inaccessible to pooled analyses. Systematic sgRNA-ablation experiments further showed that densely tiled designs improve event recovery and positional stability, providing concrete experimental guidance for future genomic locus proteomics studies.

## 1 Background

Proximity proteomics has revolutionized protein interactome research, beginning with the BirA-based BioID system [1] and rapidly expanding to include kinetically faster enzymes such as TurboID and APEX2 [2]. Several groups have fused these enzymes to nuclease-dead Cas9 (dCas9) to anchor proximity labeling at user-defined genomic loci and identify proteins associated with the specific region, an approach known as genomic locus proteomics. Established implementations include CASPEX (dCas9–APEX2) [3], C-BERST [4], and CasID [5]. Each uses one or more sgRNAs to anchor a biotinylation enzyme to the target genomic locus, followed by mass spectrometry-based identification of the labeled proteome.

These experiments routinely identify dozens to hundreds of proteins enriched at a chromatin locus. However, the analytical layer that turns those protein lists into testable transcription-factor and chromatin-regulator predictions has remained underdeveloped. Per-region logFC values are typically pooled across guides for hit calling [3, 6, 7], discarding the spatial information that multi-guide tiling could in principle provide. Downstream analysis then usually involves manually cross-referencing hits against motif databases such as JASPAR [8] or HOCOMOCO [9, 10] and against ChIP-Atlas [11] for orthogonal binding support, with no systematic deconvolution of where within the labeled window each factor likely binds.

To enable automated spatial reconstruction of genomic locus proteomics experiments, here we describe GLproxScape (**G**enomic **L**ocus **prox**imity proteomics land**scape** analysis), an R package that takes per-region proximity proteomics enrichment values as input and applies a Gaussian labeling-kernel forward model to produce motif-anchored binding-event predictions for transcription factors and zone-based binding predictions for chromatin regulators, with per-complex member co-localization. We demonstrate GLproxScape on diverse datasets [3, 6, 7, 12] and show that the same labeling-kernel model recovers known regulatory complexes at each locus and adds per-zone spatial detail that pooled-condition analyses cannot reveal.

## 2 Results

### 2.1 GLproxScape predicts region-specific transcription factor binding events

A typical genomic locus proteomics experiment produces overlapping spatial measurements in which each sgRNA samples a partially distinct chromatin neighborhood through proximity labeling (Fig. 1A). GLproxScape takes these per-region tables together with an sgRNA-to-region manifest (Fig. 1B), classifies each protein against curated transcription-factor and chromatin-regulator lists, and routes them through three analytical paths: (i) motif-anchored deconvolution for transcription factors with a motif entry, (ii) motif-free peak prediction for transcription factors absent from the motif database, and (iii) zone-based detection for chromatin regulators that lack sequence-specific motifs.

**Figure 1:**
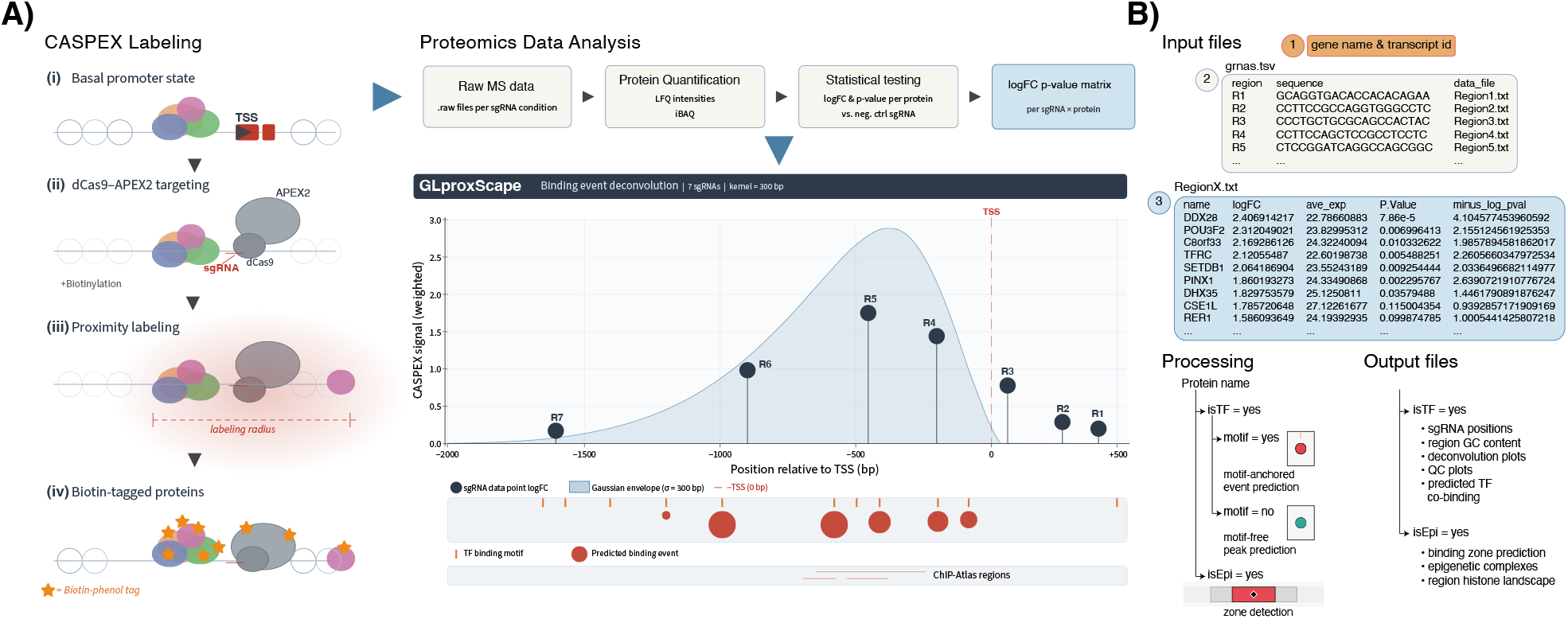
GLproxScape converts tiled genomic locus proteomics data into spatial chromatin binding predictions. A) Experimental workflow: CASPEX (dCas9–APEX2) is targeted to a chromatin region by multiple tiled sgRNAs; biotinylated proteins are captured, identified by mass spectrometry, and per-region differential statistics are computed. GLproxScape then converts these enrichment tables into spatially resolved transcription-factor binding events (motif-anchored or motif-free) and chromatin-regulator binding zones. B) Pipeline schematic: required inputs (gene name and transcript id, sgRNA-to-region manifest, per-region differential statistics), internal processing logic and generated outputs.

To evaluate the performance of GLproxScape, we first reanalyzed the dataset of Myers et al. [3] at the human TERT and MYC promoters (TERT shown here; MYC results are available on Zenodo [13]). Per-region logFC correlation showed strong agreement across R2–R5 with R1 separating from the rest (Fig. S1A), consistent with the distal placement of R1 relative to the tightly clustered R2–R5 design. Motif-driven deconvolution recovered binding events for MAZ, ZXDC, ZIC3, and FOXP2 (Figs. 2A and S1B), and for MAZ specifically 7 of the 12 predicted events overlapped with ChIP-Atlas peaks (in ± 25bp distance range) from three independent experiments (Figs. 2A and 2C), providing orthogonal support. Consistent with Myers et al. [3], GLproxScape recovered strong enrichment of zinc finger transcription factors (ZNFs) at the TERT promoter (Fig. S1C): six ZNFs with JASPAR motifs (ZNF16, ZNF76, ZNF148, ZNF184, ZNF214, ZBTB24) received motif-anchored binding-event predictions (Fig. 2B), while additional ZNFs without JASPAR entries were resolved by motif-free peak prediction (Fig. S1D). In addition to ZNFs, GLproxScape predicted MAP3K7 binding to TERT locus, which was among the several confirmed hits in the original paper [3] (Fig. S1E).

**Figure 2:**
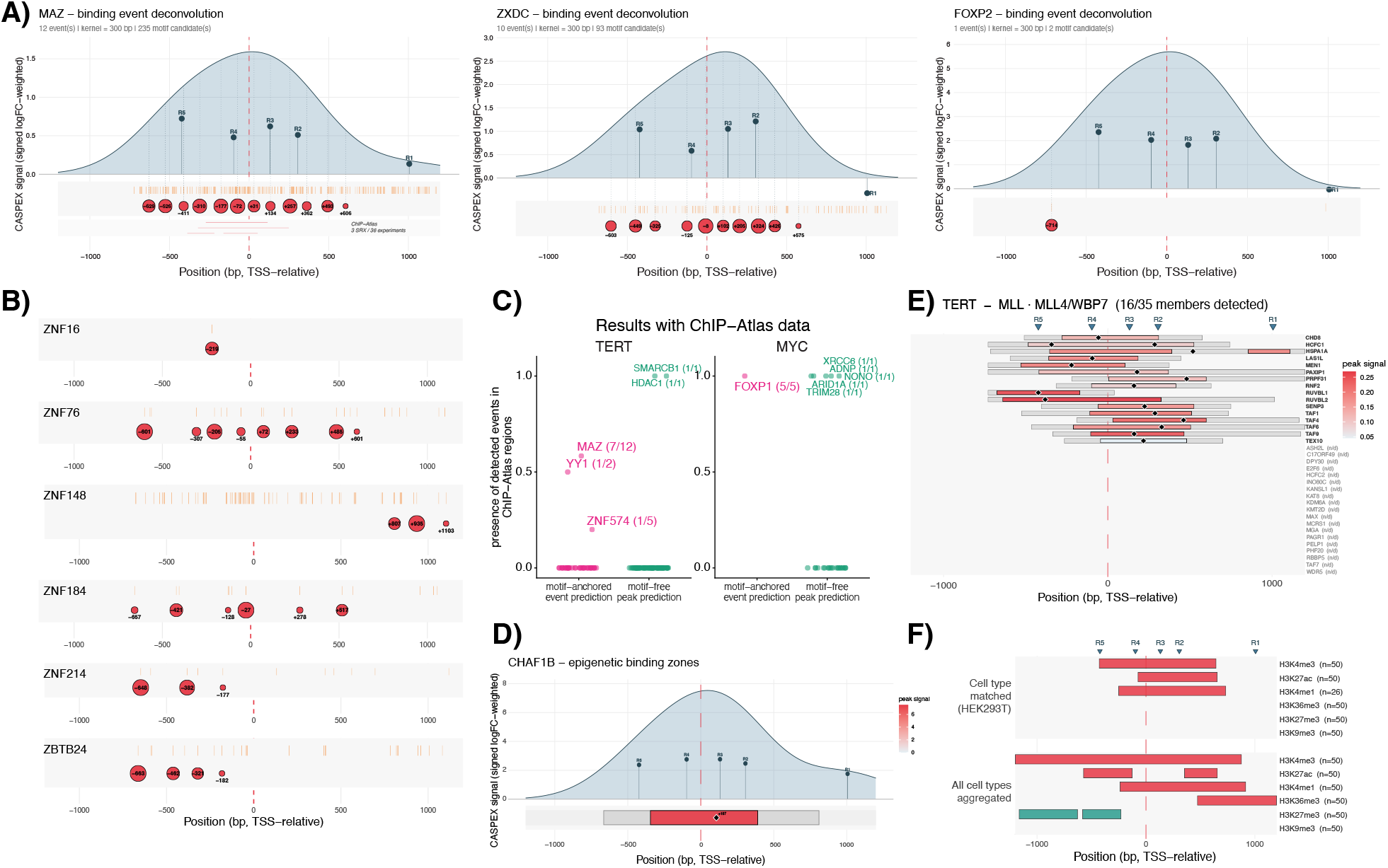
GLproxScape analysis of TERT chromatin region genomic locus proteomics enrichments. A) Analysis of GLproxScape results and motif-anchored binding predictions for MAZ, ZXDC, and FOXP2 in TERT genomic region. B) Motif-anchored binding predictions for ZNF16, ZNF76, ZNF148, ZNF184, ZNF214 and ZBTB24 in TERT genomic region. C) The coverage ratio of predicted events in ChIP-Atlas data (matched within ± 25 bp) in TERT and MYC datasets. D) Epigenetic binding zone prediction of CHAF1B on TERT genomic region. E) Predicted binding zones of MLL4/WBP7 complex on TERT genomic region. F) GLproxScape analysis of histone marks on TERT genomic region.

Scanning predicted binding events for co-localization within ± 50 bp revealed (i) recurrent FOXP2– FOXP4–FOXK2 co-binding consistent with their shared motif preferences [14], (ii) broad co-localization of ETV3, MAZ, and NR2F1 with multiple other factors, and (iii) ZXDC co-localization with many ZNFs (Fig. S1F). Comparison against ChIP-Atlas datasets showed that several proteins such as MAZ (7/12), YY1 (1/2), and FOXP1 (5/5) had at least half of their predicted binding events supported by overlapping ChIP-Atlas peaks in the TERT and MYC datasets (Fig. 2C).

We additionally tested how kernel-width and coverage-floor choices affect predicted event positions. Across the *σ* sweep, most TF predictions were stable at *σ* ≥300 bp (e.g., ZBTB24, FOXP2), while MAZ event positions drifted at *σ* < 300 bp before stabilizing (Fig. S1G), consistent with short G-rich motif of MAZ [15] producing densely-spaced PWM hits at the GC-rich TERT promoter [16], where narrower kernels fail to resolve adjacent candidate positions into stable peaks [17]. Predicted event positions were robust to coverage-floor settings between 0.03 and 0.15 (Fig. S1H).

Together, these results show that GLproxScape recovers both motif-anchored and motif-free TF binding events from a genomic locus proteomics dataset, validates a subset against publicly available ChIP-Atlas data, and reveals candidate co-binding pairs.

### 2.2 GLproxScape identifies possible epigenetic factor binding zones

Epigenetic regulators, including chromatin readers, writers, erasers, and remodelers typically lack a sequence-specific DNA-binding motif, instead, they recognize chemical changes on the DNA and histones. To address this difference, GLproxScape performs zone prediction for epigenetic factors. Analysis of TERT genomic region predicted a high CHAF1B enrichment through R2–R5, which was validated with targeted ChIP-qPCR experiments in Myers et al. [3] (Fig. 2D). An epigenetic complex focused analysis revealed the detection of MLL4/WBP7 complex (16 of 35 members) at this region with overlapping predicted zones (Fig. 2E). Additionally, GLproxScape performs enrichment for key histone modifications to allow the direct comparison of predicted zones with the histone modification landscape of the region (Fig. 2F).

### 2.3 GLproxScape robustly performs on different datasets and species

To evaluate GLproxScape on another dataset, we reanalyzed the FOXP2 genomic locus of MacKenzie et al. [6] (3 sgRNAs spanning 2 regions). GLproxScape recovered strong motif-anchored predictions for MAX and ETV6 (Figs. 3A and S2C), with 3/6 and 7/7 of their predicted events, respectively, overlapping peaks from independent ChIP-Atlas experiments. The strongest FOXA1 motif-driven prediction was supported by 62 independent ChIP-Atlas experiments despite only modest logFC enrichment (Fig. 3B, S2C), showing that GLproxScape surfaces biologically plausible events even when the proteomic signal is weak. Top predicted events for GATA4 and TFAP4 likewise overlapped with ChIP-Atlas peaks (Figs. 3C and S2C), as did the motif-free HMGB1 prediction (Figs. 3D and S2C). Predicted events showed dense TF co-localization within ± 50 bp (Fig. S2A). Distribution of TF predictions showed that ETV6 and SALL1 are the factors with top region-specific enrichment, while ZNF112, ZNF184, ZNF286B, and ZFHX2 showed high composite enrichment (Fig. S2B). Among these, ETV6, GATA4, and ZFHX2 were reported as significantly enriched by the original authors, with ZFHX2 further validated by ENCODE ChIP-Seq [6].

**Figure 3:**
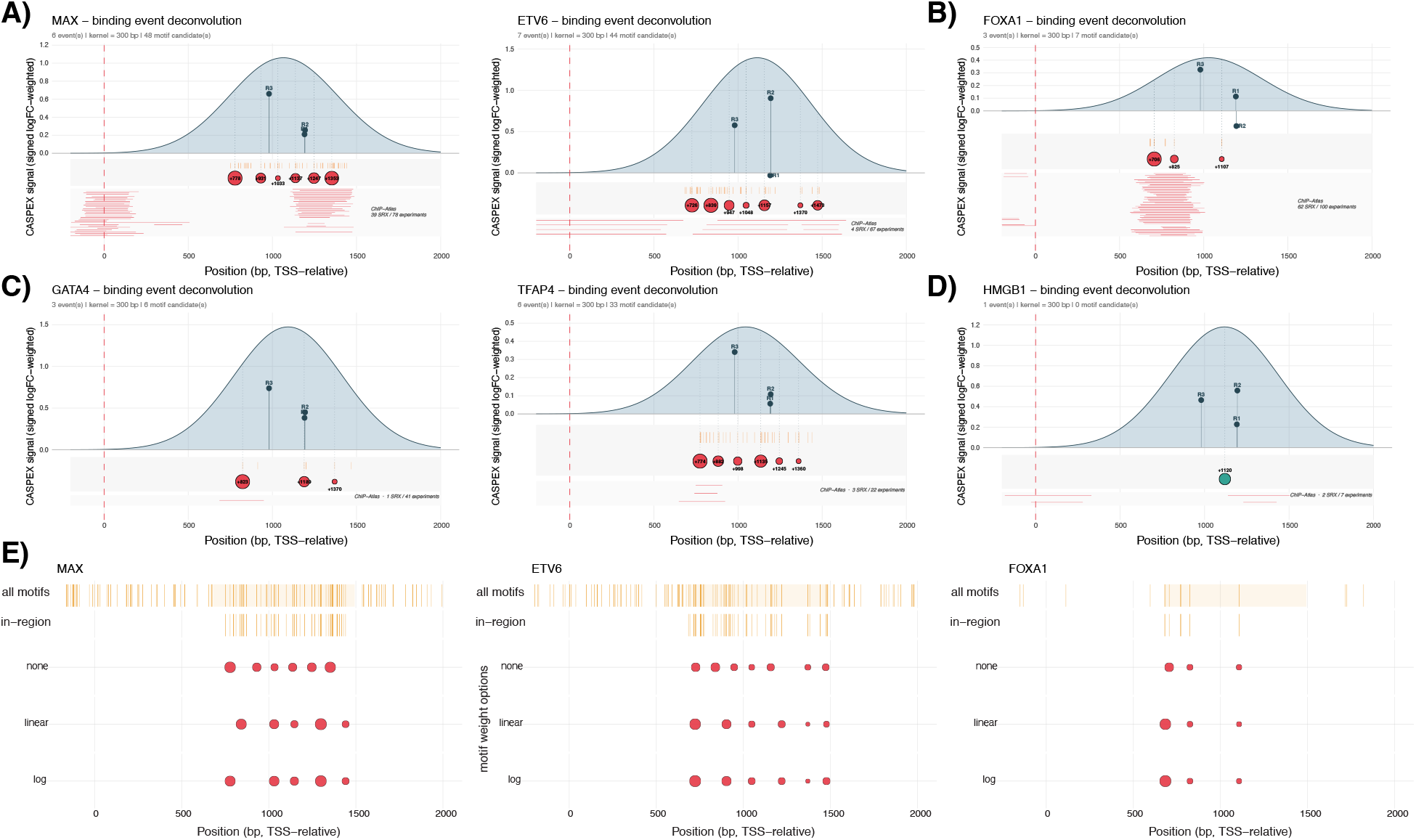
GLproxScape analysis of FOXP2 genomic region data. A-D) Predicted binding events of MAX and ETV6 (A), FOXA1 (B), GATA4 and TFAP4 (C), and HMGB1 (D) on FOXP2 genomic region. E) Predicted binding site layouts with motif weight options (linear or logarithmic) for representative TFs (ETV6, MAX, FOXA1). Predicted event bubble sizes may appear different from those in other panels because a shared absolute scaling was applied to enable direct comparison between three conditions.

GLproxScape analyses of the TERT, MYC, and FOXP2 datasets used logFC-driven signal weighting without rescaling the binding predictions by JASPAR PWM scores. Applying linear or logarithmic PWM weighting produced only subtle changes for ETV6, MAX, and FOXA1 (Fig. 3E). The convergence across weighting schemes argues that calls are driven by the proximity-labeling signal, not by PWM-score choices; we therefore use unweighted scoring as the default.

Beyond PWM weighting, we also tested whether the choice of motif database itself affects GLproxScape predictions by re-running the FOXP2 analysis with HOCOMOCO v12 [9, 10] as an alternative position-weight-matrix source. Of the FOXP2 spatial-model TFs with a motif in either database, 23 were shared between JASPAR and HOCOMOCO, 16 were JASPAR-only, and 4 were HOCOMOCO-only (Fig. S2D). For the top eight TFs by combined NNLS amplitude across both engines, predicted event positions tracked closely between the two databases (Fig. S2E), events called by both engines validated against ChIP-Atlas at a higher rate than events called by either engine alone (Fig. S2F), and this concordance was preserved across all three JASPAR weighting variants (Fig. S2G). An unbounded nearest-neighbour analysis confirmed this concordance quantitatively: 62–70% of motif-anchored events had a same-TF call in the other database within ±50 bp, rising to 83–89% within ±100 bp (Fig. S2H), with per-TF match rates at ±50 bp ranging from near-complete concordance (≥75% in both directions for KLF3, MEIS1, ZEB2, MEIS3, ETV3) to weaker agreement for TFs with divergent matrix representations between the two libraries (e.g. SOX2, SIX1; Fig. S2I). Together, these results indicate that spatial predictions of GLproxScape are driven primarily by the proximity-labeling signal rather than the motif source or scoring scheme.

Additionally, we tested GLproxScape on a mouse cell line dataset [12] and recovered motif-anchored binding events for NFKB1 and NFKB2 at the Ripk3 promoter; NFKB1 was independently validated in the original paper, while the other validated regulator IKBKG is absent from the reference TF library, but extending the TF universe to include it recovered IKBKG via the motif-free peak-prediction path (Figs. S3A and S3C), demonstrating that user-defined TF universes can accommodate regulators that fall outside canonical sequence-specific TF lists. A nearest-neighbour analysis between JASPAR and HOCOMOCO on this dataset further showed strong cross-database agreement (~70% match rate at ± 50 bp in both directions; Fig. S3B). Re-running the same dataset with the mouse-specific HOCOMOCO PWM bundle in place of vertebrate-pooled JASPAR matrices recovered substantially more NFKB1 events at the Ripk3 promoter (1→ 4 events) with higher event amplitudes (max weight 3.5 →13.9; Fig. S3C), suggesting that species-matched motif libraries can meaningfully improve sensitivity.

### 2.4 Dense sgRNA tiling and *p*-value weighting enhance spatial prediction sensitivity

To evaluate the *p*-value-driven weighting mode, we first analyzed a published dataset targeting the FOXQ1 promoter with four densely-tiled sgRNAs and two biological replicates per guide [7]. GLproxScape predicted motif-anchored binding events for FOXC1, RBPJ, and SP1 at the FOXQ1 locus, all supported by overlapping ChIP-Atlas peaks (Fig. S4A). MAZ was also recovered with multiple events across regions (Fig. S4B). Notably, *p*-value-driven weighting additionally recovered binding predictions even when the per-region enrichment pattern was strongly imbalanced, as for RREB1, where only a single sgRNA (R2) carried positive signal and the other three regions were near zero (Fig. S4C), and for KLF16 in a more extreme case where three regions showed explicitly negative enrichment with only one sgRNA contributing positive signal (Fig. S4D). Under logFC-driven weighting, such asymmetric per-region patterns would attenuate the per-protein composite and suppress the call; *p*-value weighting instead retains the spatially localized signal carried by the surviving region.

Next, we tested whether *p*-value-driven weighting also performs well on more sparsely tiled sgRNA designs spanning a larger genomic window. We analyzed an in-house dataset targeting a ~3000 bp locus with 7 tiled sgRNAs and two biological replicates per region. We specifically examined TFs with modest logFC enrichment to test whether *p*-value-based weighting could recover spatially localized regulatory signals that would be under-emphasized by logFC-driven analysis alone. GLproxScape identified a motif-free binding-zone prediction for MTA2 that overlapped with independent ChIP-Atlas peaks (Fig. 4A). In addition, the strongest motif-anchored prediction for ZNF740 localized near the transcription start site and was supported by ChIP-Atlas data despite the absence of strong local logFC enrichment in the corresponding region (Fig. 4B). Notably, this prediction was enabled by signal contribution from region 3, illustrating how tiled sgRNA designs can improve spatial sensitivity by capturing distributed enrichment patterns across neighboring regions.

**Figure 4:**
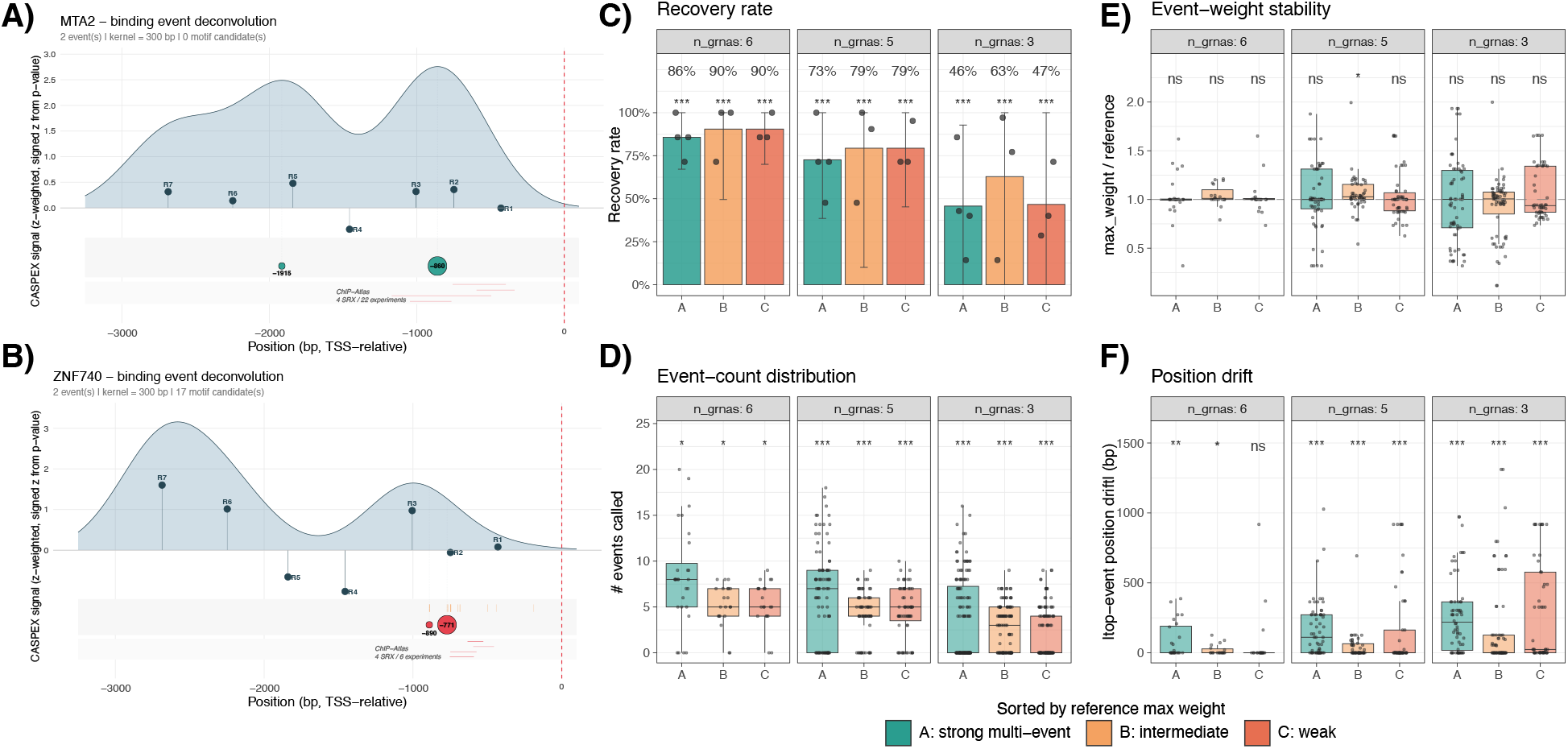
GLproxScape successfully identifies binding sites with *p*-value weighting option and dense gRNA tiling. A) Motif-free peak prediction of MTA2 on a promoter region. B) Motif-anchored prediction of ZNF740 on a promoter region. C-F) Region (gRNA) ablation simulations show progressive deterioration of spatial prediction with reduced gRNA number on the promoter. Starting from the full 7-gRNA reference, all C(7,6)+C(7,5)+C(7,3) = 63 subsets were re-run through GLproxScape and compared to the reference reconstruction for the top-10 motif-anchored TFs, grouped into tiers A/B/C (strong multi-event / intermediate / weak single-event) by reference weight tertile× event count. C) Recovery rate per tier; bars show the mean per-TF recovery with 95% *t*-CIs across TFs, stars from a one-sided exact binomial test against the n=7 reference. D) Per-(TF, subset) event counts; a paired Wilcoxon signed-rank test was applied. E) Event-weight stability of the top event per TF; a one-sample Wilcoxon signed-rank test was applied. F) Absolute positional drift in bp of the top event relative to the 7-gRNA reference; a one-sample Wilcoxon signed-rank test was applied. All *p*-values BH-adjusted within each metric across the nine (tier × dropout-size) cells. Significance: * *q* < 0.05, ** *q* < 0.01, *** *q* < 0.001, ns: not significant.

Given that tiled sgRNA density is central to the spatial reconstruction framework, we next evaluated prediction robustness under systematic guide ablation. Starting from the full 7-guide dataset, we generated all lower-density combinations of 6-, 5-, and 3-guide configurations and compared the recovered predictions against the full-reference reconstruction (Figs. 4C–F). Reduced guide number progressively decreased both the recovery rate and the total number of recovered events (Figs. 4C and 4D), supporting the importance of dense tiling for stable spatial inference. In contrast, no significant global effect was observed on predicted event weights across ablation groups (Fig. 4E), indicating that guide loss primarily affects event detectability and positional stability rather than the inferred strength of retained predictions. Guide ablation additionally introduced positional drift relative to the 7-guide reference reconstruction, with TFs with weak predictions showing greater displacement than strong predictions (Fig. 4F).

## 3 Discussion

Tiled genomic locus proteomics experiments intrinsically generate overlapping spatial measurements because each sgRNA samples a partially distinct chromatin neighborhood through a finite labeling radius. However, conventional analyses primarily used multiple sgRNAs to reduce the background noise [3], which collapses spatial structure by pooling enrichment values across regions before hit calling [6, 7], reducing the experiment to a genomic locus level protein list. GLproxScape instead treats observed enrichments as convolutions of an underlying latent occupancy landscape and models each sgRNA as a continuous Gaussian biotinylation source. This enables the reconstruction of coverage-normalized spatial enrichment profiles and converts tiled proteomics measurements into positional regulatory predictions. The framework further separates predictions into three analytical paths: (i) motif-anchored NNLS deconvolution for sequence-specific TFs with PWM entries, (ii) motif-free peak prediction for TFs absent from the PWM database, and (iii) zone-based reconstruction for broad chromatin-associated factors. This distinction is important because forcing epigenetic regulators into point-wise TF-style predictions would misrepresent their occupancy patterns.

Across reanalyzed datasets, preserving spatial resolution enabled several forms of biological interpretation that are inaccessible in pooled analyses. First, motif-resolved event positioning enabled local predicted co-binding analysis, recovering recurrent TF associations such as FOXP2–FOXP4–FOXK2 and ZXDC co-localization with multiple ZNFs at the TERT locus. Second, spatial overlap between predicted chromatin-factor zones allowed reconstruction of higher-order regulatory assemblies, including partial recovery of the MLL4/WBP7 complex with overlapping inferred occupancy domains. Third, the framework recovered biologically plausible regulatory events even under weak enrichment conditions. For example, the FOXA1 prediction at the FOXP2 locus showed extensive orthogonal support from ChIP-Atlas despite only modest local enrichment, indicating that biologically relevant recruitment patterns may remain detectable after spatial reconstruction even when conventional enrichment ranking would deprioritize them. Fourth, the spatial reconstruction strategy generalized across species, as re-running the framework on the mouse Ripk3 dataset with species-matched motif (HOCOMOCO v12 mouse) and assembly-aware ChIP-seq (mm10) references recovered the validated regulators (NFKB1 and IKBKG), suggesting that the analytical strategy is not tied to human-specific reference panels.

The analyses also provide practical guidance for future experimental design strategies. Dense sgRNA tiling substantially improved spatial robustness, while guide ablation progressively reduced event recovery and increased the positional instability. Importantly, reduced guide number affected event detectability more strongly than recovered event amplitude, suggesting that sparse tiling primarily limits localization accuracy and reconstruction completeness rather than signal magnitude itself. The increased positional drift observed among weakly supported TF predictions further indicates that low-confidence calls are disproportionately sensitive to guide sparsity. Together, these findings support the use of densely tiled designs for stable spatial inference in genomic locus proteomics experiments. In parallel, datasets with biological replicates and calibrated per-region statistics benefited from *p*-value-driven weighting, which enabled recovery of spatially localized events that would likely be overlooked by logFC-based weighting alone.

Several limitations should be considered when interpreting GLproxScape predictions. First, the forward model currently operates strictly along the linear genome and does not account for three-dimensional chromatin organization. As a consequence, proteins recruited to a target region through long-range looping contacts will be assigned to linear positions, and may therefore appear as false-localized events. Second, the motif-resolved path depends on the completeness of available PWM databases, particularly JASPAR, whose coverage remains uneven for non-canonical or poorly characterized DNA-binding proteins. The motif-free fallback provides approximate spatial localization for these factors, but remains heuristic and does not impose a sequence-based mechanistic constraint. This caveat is partially mitigated by the database-agnostic structure of the motif-resolved path itself, as re-running the FOXP2 analysis with HOCOMOCO v12 produced near-identical event positions on shared TFs (Figs. S2D–G), indicating that database choice modulates which TFs are recoverable rather than where their events are placed. Relatedly, the choice of PWM weighting scheme (unweighted, linear, or log) materially affects which candidates surface from the deconvolution; given that JASPAR scores carry real affinity information, hits that emerge only under linear or log weighting may still reflect biologically meaningful binding rather than a methodological artifact, and we recommend cross-checking results across weighting modes. Third, orthogonal validation against ChIP-Atlas is itself imperfect, reflecting uneven antibody availability, variable cell-type representation, and heterogeneous ChIP-seq quality across experiments. Overlap with ChIP-Atlas peaks therefore constitutes supporting evidence for GLproxScape predictions rather than a formal estimate of accuracy. Importantly, ChIP-Atlas is an actively curated resource that continues to expand with additional ChIP-seq experiments, broader antibody coverage, and more diverse cell-type representation; as the compendium grows, its capacity to corroborate GLproxScape predictions is expected to strengthen accordingly, and predictions that currently lack orthogonal support, particularly for sparsely profiled factors or under-represented cellular contexts, may gain confirming evidence as the reference dataset matures, refining the apparent precision of the framework without any change to the underlying model.

The framework nevertheless provides several opportunities for future extension. Incorporating chromatin conformation information could relax the assumption of purely linear genomic distance and improve the interpretation of distal recruitment events. The kernel-based forward model is additionally agnostic to the specific proximity-labeling enzyme and should generalize to alternative systems such as TurboID or miniTurbo after kernel recalibration. Finally, because the reconstruction framework depends primarily on tiled spatial sampling rather than promoter-specific assumptions, the same analytical strategy should generalize to enhancers, super-enhancers, and other regulatory genomic elements.

Together, these analyses establish GLproxScape as a general spatial-reconstruction framework for tiled genomic locus proteomics. The strategy generalized across human and mouse datasets with species-matched motif and ChIP-seq references, and across motif databases and weighting schemes without changes to the underlying engine. Because the forward model relies on tiled spatial sampling rather than promoter-specific assumptions, the same approach is positioned to extend to enhancers and alternative proximity-labeling enzymes.

To our knowledge, the four studies reanalyzed here [3, 6, 7, 12] constitute most currently available genomic locus proteomics datasets that employed tiled sgRNA designs, the input format required for spatial reconstruction. As published genomic locus proteomics compendia accumulate, data-driven refinements such as learning per-TF positional priors or labeling-kernel parameters could further complement the framework. We anticipate that GLproxScape will provide a useful computational backbone for the next generation of genomic locus proteomics studies.

## 4 Methods

### 4.1 Inputs and per-guide signal model

GLproxScape takes two inputs per dataset: a tab-separated sgRNA manifest mapping each region label to its 20-bp protospacer sequence and per-region differential file, and one differential-statistics file per region containing protein name, log fold-change, and *p*-value columns. The pipeline does not perform peptide-to-protein assembly or differential testing itself; users supply these from any standard proteomics pipeline (e.g. `limma` [18], MSstats [19], DEP [20], or a custom statistical procedure).

Per-region per-protein signal is encoded as a signed *z*-score derived from the *p*-value by default, as a moderated *t*-statistic when one is available from the differential pipeline (e.g. `limma`), or as a signed log fold-change when *p*-values are saturated or under-powered. Promoter sequence and gene/transcript coordinates are retrieved from Ensembl via REST API.

Before signal construction, each protein passes a per-region significance gate: detected in at least `min_regions` regions (default 2) at *p* ≤ `pval_thresh` (default 0.05, relaxed where per-region *p*-values are uninformative), with an optional `min_lfc` floor on positive enrichment. Proteins clearing these gates enter the spatial model.

The biotinylation radius of APEX2 fused to nuclease-dead Cas9 sets a soft labeling profile around each guide’s binding site. Following the labeling-radius estimate of Hung et al. [2] (~10–20 nm in 3D, corresponding approximately to a few hundred base pairs along linear DNA), GLproxScape models the labeling profile of guide *r* centered at promoter position *g*_*r*_ using a Gaussian with *σ* = 300 bp by default. For each protein *p*, the per-region enrichment weight 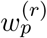 (signed *z*-score, moderated *t*-statistic, or signed logFC, selected by `weight_mode`) is forward-smeared into a continuous spatial track:

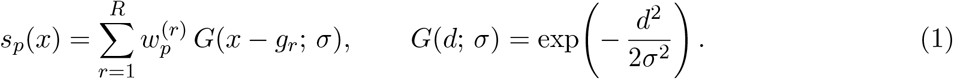

The joint coverage profile *C*(*x*) = ∑_*r*_ *G*(*x* − *g*_*r*_; *σ*) summarizes how reachable each position is from *some* sgRNA, and a coverage-normalized signal

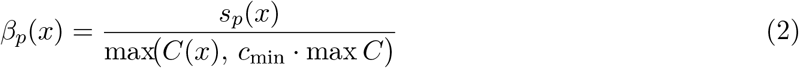

is computed with floor *c*_min_ = 0.05 (`cov_floor`) to suppress division blow-up at the gRNA-tile edges where *C*(*x*) drops toward zero. The forward smear encodes the labeling-radius assumption that high 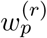 contributes a larger local labeling cloud near *g*_r_, pulling the predicted binding peak toward high 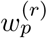 regions; the coverage normalization removes the bias toward dense-tiling interiors that arises when the same protein is sampled by overlapping guides.

### 4.2 Motif-resolved binding-event deconvolution for transcription factors

For each transcription factor *τ* present in the user-supplied TF list (default: a TF list derived from Lambert et al. [21]) with a JASPAR entry [8], GLproxScape retrieves the position weight matrix *M*_*τ*_, log-odds scores it under a uniform background (*p* = 0.25 per base) with pseudocount 0.25, and scans the promoter sequence at a threshold of 0.80*·* maxScore as default. Motif hits at positions {*m*_*τ,i*_} _*i*_ define a non-negative basis whose amplitudes *β*_*τ,i*_ ≥ 0 are recovered by non-negative least squares (NNLS):

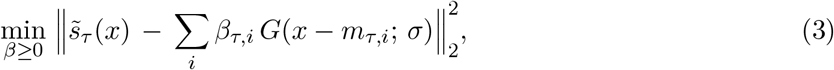

where 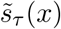 is the coverage-normalized spatial track for TF *τ*. By default, every motif hit enters this basis with unit weight, so *β*_*τ,i*_ depends only on the alignment of 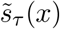 with each Gaussian centered at *m*_*τ,i*_. GLproxScape optionally rescales basis columns by a function of the per-hit PWM-score fraction *f*_*τ,i*_ = (*S*_*τ,i*_ − *S*_min_)*/*(*S*_max_ − *S*_min_) ∈ [`threshold_frac`, 1], where *S*_*τ,i*_ is the raw log-odds score of hit *i* and *S*_min */* max_ are the matrix extremes: `motif_score_weight = “linear”` sets the column scale to *f*_*τ,i*_, `“log”` to 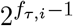, and `“none”` (default) leaves the basis unscaled. Score weighting biases recovered *β* toward higher-scoring motifs within clusters of nearby hits.

Two boundary controls trim deconvolution events near the gRNA-tile edges where the coverage prior is no longer trustworthy. First, *β* is masked to the in-support region *C*(*x*) > `edge_guard_frac` *·* max *C* (default 0.25, i.e. 5× the `cov_floor` clamp) so the floor-induced ramp at the tile boundaries lives outside the deconvolution domain. Second, events whose distance to the nearest gRNA exceeds `max_grna_distance` (default *σ*, scaling naturally with the kernel width) are dropped, preventing the Gaussian tail of a single boundary guide from anchoring a call far from any actual labeling source.

Events with *β*_*τ,i*_ below `min_weight_frac` (default 0.15) of the local peak amplitude are pruned. Surviving events within `merge_dist` bp (default 100 bp; *≈σ/*3 at *σ* = 300 bp) are merged into amplitude-weighted clusters; the cluster’s reported position is snapped to the strongest motif in the cluster (`merge_position = “argmax”`). A final cap of `max_events_per_tf` (default 30) retains the top events per TF by combined cluster amplitude, preventing unreadable plots on promiscuous PWMs with broadly elevated signal. TFs detected in the proteomics but absent from JASPAR fall through to a motif-free peak-detection branch that emits one event at the spatial track’s argmax under the same coverage prior. Each surviving event is reported with its TSS-relative position, fitted weight, and supporting motif-hit count.

While JASPAR is the default motif source, GLproxScape also supports HOCOMOCO v12 [9, 10] as an alternative position-weight-matrix database via the `motif_search_engine` argument; the spatial-deconvolution machinery downstream of the motif scan is database-agnostic.

### 4.3 Zone-based deconvolution for chromatin factors

Epigenetic regulators typically lack sequence-specific DNA-binding motifs and bind in domains rather than at points; the motif-NNLS path is therefore inappropriate for these factors. For each factor in a user-supplied epigenetic-factor list (default: EpiFactors database [22]), GLproxScape’s epigenetic deck replaces the motif basis with a zone-based deconvolution that operates directly on *s*(*x*) rather than *β*(*x*).

#### Zone detection

The spatial track *s*(*x*) is thresholded at zone_frac *·* max(*s*) (default `zone_frac` = 0.30); the contiguous bp range above the threshold is reported as the *outer zone*, representing the broad chromatin-domain extent. The choice `zone_frac` = 0.30 produces ~930 bp baseline zones for single-source signals at *σ* = 300 bp, consistent with the kernel resolution limit, while merging multi-source plateaus into broader domains. GLproxScape analyses use *s*(*x*) rather than *β*(*x*) because for factors with relatively uniform per-region logFCs (typical of broad-binding chromatin readers) *β* = *s/C* amplifies at lone boundary guides where both numerator and denominator collapse to a single guide’s contribution, putting the apparent peak at the gRNA-tile edge even when labeling intensity is genuinely highest in the multi-guide interior. *s*(*x*) directly represents labeling intensity, the quantity proximity labeling actually measures.

#### Multi-core focal detection

Within each outer zone, all contiguous supra-threshold runs of *s*(*x*) > inner_zone_frac*·* zone_max (default 0.70) are detected and rendered as darker inner-core bars with peak-signal-colored fill. A zone with one sharp peak yields one core; a zone with two well-separated sub-peaks separated by a sub-threshold valley yields two cores; etc. This preserves the kernel-honest broad-domain framing of the outer zone while marking where predicted binding is most concentrated.

#### Region-specific centroids

For each focal sub-peak, candidate centroids are gRNA regions inside the zone that are (i) local logFC maxima and (ii) carry at least `centroid_frac` (default 0.70) of the zone’s maximum regional logFC. Each centroid’s bp coordinate is then computed as the logFC-weighted average of its region and the immediate positive-logFC neighbours. Centroids are calculated coordinates, not gRNA-snapped, so a zone with two high-logFC clusters separated by lows can yield two centroids each pulled toward its supporting cluster. Centroid positions may differ from *s*(*x*)’s argmax because kernel-summed peaks can be biased toward multi-guide overlap regions even when one region carries the standout individual signal.

### 4.4 Validation against ChIP-Atlas

For each transcription factor and chromatin regulator in the deck, GLproxScape queries ChIP-Atlas [11] for matching ChIP-seq peak BED files (per-SRX, threshold *Q* < 1× 10^−5^ by default) and renders peaks falling within the analysis window as a stacked sub-lane below the predicted binding-event bubbles. The reference assembly is set by `chipatlas_genome` (default hg38) and is auto-derived from the `species` argument, so cross-species runs require only a single species declaration rather than separate per-component genome settings. When the source coordinates differ from the ChIP-Atlas assembly, as for mouse runs where Ensembl serves GRCm39 while ChIP-Atlas remains on mm10, the analysis window is lifted over to the ChIP-Atlas assembly via the Ensembl REST archive before peak overlay, with lift-over results cached locally to avoid repeated remote calls.

The default cap is the 100 most recent SRX per factor (ordered by descending accession) and can be raised globally. For biologically focal regulators where cell-type-specific binding may live in older SRX entries that the cap excludes, the `special_interest_gene` parameter accepts a character vector of gene symbols whose scan bypasses the standard cap; an optional `special_interest_cap` sets a separate (typically larger) ceiling for these factors, defaulting to all available SRX. Per-SRX BED files are cached locally under R’s user-cache directory so cross-locus and cross-dataset analyses share downloads. The deck annotation reports the number of SRX with peaks in the analysis window over the number of SRX scanned (e.g., `8 SRX / 10 experiments`); special-interest factors are marked by a trailing asterisk.

### 4.5 Locus-level histone-mark landscape

In addition to the per-factor ChIP-Atlas overlay, GLproxScape writes a single `histone_marks.pdf` per locus summarizing the chromatin-state landscape, with two stacked sections of six rows each. The top section shows the six canonical marks (*active*: H3K4me3, H3K27ac, H3K4me1, H3K36me3; *repressive*: H3K27me3, H3K9me3) restricted to ChIP-Atlas SRXs whose `cell_type` field matches the user-supplied experimental cell line (`histone_cell_type`) by case- and punctuation-insensitive substring, capped at `histone_max_experiments_matched` (default 50). The bottom section repeats the same six marks aggregated across all cell types in ChIP-Atlas, capped at `histone_max_experiments_all` (default 50). Each row renders the union of peaks across qualifying SRXs as horizontal bars, with active marks colored red and repressive marks teal so the chromatin state reads at a glance. Empty rows still draw with their label and SRX-count annotation so absence-of-data is visually distinct from absence-of-signal.

### 4.6 Non-default run parameters for different datasets

For the TERT and MYC datasets [3], we used signed-logFC weighting because the manuscript does not report per-region *p*-values. For the FOXP2 dataset [6], we kept signed-logFC weighting because its per-region contrasts are statistically under-powered relative to the pooled (*n* = 9) approach used in the original analysis. For the FOXQ1 dataset [7], the Ripk3 dataset [12], and the in-house dataset, all of which include biological replicates with calibrated per-region statistics, we used the *p*-value-driven (signed-*z*) weighting mode.

For the mouse Ripk3 dataset, the `species` argument is additionally set to mouse, which auto-derives the HOCOMOCO mouse PWM bundle, the mm10 ChIP-Atlas reference, and the GRCm39 *→*mm10 lift-over described in §4.4. The TF universe is also extended to include IKBKG, the second regulator validated in the original paper, so that it is evaluated via the motif-free peak-prediction branch despite falling outside the canonical sequence-specific TF list. Across all datasets, the promoter window is set per dataset based on the sgRNA tile span, and transcript anchors are pinned to a non-canonical Ensembl transcript when the canonical one does not match the experimentally targeted promoter. Exact parameter values for each run are recorded in the dataset runner scripts deposited in the Zenodo repository [13].

## Data availability

All analysis input files, preprocessing files, and result outputs generated in this study have been deposited in Zenodo [13]. The in-house dataset is associated with a separate study currently in preparation; the corresponding citation information will be added upon publication. A detailed comparison of GLproxScape results with the original papers is available in Supplemental Table - 1.

## Code availability

All analysis scripts have been deposited in the Zenodo repository. The GLproxScape R package can be directly installed from GitHub (https://github.com/scanozcan/GLproxScape).

## Acknowledgments

The authors gratefully acknowledge the use of the services and facilities of the Koç University Research Center for Translational Medicine (KUTTAM), funded by the Presidency of Turkey, Presidency of Strategy and Budget. We thank Dr. Stefan Koch for sharing the FOXQ1 genomic locus proteomics sgRNA sequences and Dr. Courtney Griffin for her kind help with the Ripk3 data.

## Author contributions

Conceptualization: SCO, CA. Methodology: SCO. Software: SCO. Formal analysis: SCO. Data curation: SCO, BS, BY. Validation: BY, UC, BS, MG. Visualization: SCO. Project administration: SCO, CA. Funding acquisition: CA. Writing – original draft: SCO. Writing – review & editing: MG, CA.

## References

[1] Kyle J Roux et al. A promiscuous biotin ligase fusion protein identifies proximal and interacting proteins in mammalian cells. Journal of Cell Biology, 196(6):801–810, 2012. doi: 10.1083/jcb.201112098.

[2] Victoria Hung et al. Spatially resolved proteomic mapping in living cells with the engineered peroxidase APEX2. Nature Protocols, 11(3):456–475, 2016. doi: 10.1038/nprot.2016.018.

[3] Samuel A Myers et al. Discovery of proteins associated with a predefined genomic locus via dCas9-APEX-mediated proximity labeling. Nature Methods, 15(6):437–439, 2018. doi: 10.1038/s41592-018-0007-1.

[4] Xin D Gao et al. C-BERST: defining subnuclear proteomic landscapes at genomic elements with dCas9-APEX2. Nature Methods, 15(6):433–436, 2018. doi: 10.1038/s41592-018-0006-2.

[5] Elisabeth Schmidtmann et al. Determination of local chromatin composition by casid. Nucleus, 7(5):476–484, 2016.

[6] Tim M G MacKenzie et al. In-depth characterization of the promoter-proximal proteome of the single-copy locus FOXP2. Molecular & Cellular Proteomics, 25(4):101570, 2026. doi: 10.1016/j.mcpro.2026.101570.

[7] Giulia Pizzolato et al. The tumor suppressor p53 is a negative regulator of the carcinoma-associated transcription factor foxq1. Journal of Biological Chemistry, 300(4):107126, 2024.

[8] Jaime A Castro-Mondragon et al. JASPAR 2022: the 9th release of the open-access database of transcription factor binding profiles. Nucleic Acids Research, 50(D1):D165–D173, 2022. doi: 10.1093/nar/gkab1113.

[9] Ivan V Kulakovskiy et al. Hocomoco: towards a complete collection of transcription factor binding models for human and mouse via large-scale chip-seq analysis. Nucleic acids research, 46 (D1):D252–D259, 2018.

[10] Ilya E Vorontsov et al. Hocomoco in 2024: a rebuild of the curated collection of binding models for human and mouse transcription factors. Nucleic Acids Research, 52(D1):D154–D163, 2024.

[11] Shinya Oki et al. ChIP-Atlas: a data-mining suite powered by full integration of public ChIP-seq data. EMBO Reports, 19(12):e46255, 2018. doi: 10.15252/embr.201846255.

[12] Siqi Gao et al. Genomic locus proteomic screening identifies the nf-κb signaling pathway components nfκb1 and ikbkg as transcriptional regulators of ripk3 in endothelial cells. PloS one, 16(6):e0253519, 2021.

[13] Selahattin Can Ozcan. Glproxscape analyses, May 2026. URL 10.5281/zenodo.20337799.

[14] Sridhar Hannenhalli et al. The evolution of fox genes and their role in development and disease. Nature Reviews Genetics, 10(4):233–240, 2009.

[15] Steven A Bossone et al. Maz, a zinc finger protein, binds to c-myc and c2 gene sequences regulating transcriptional initiation and termination. Proceedings of the National Academy of Sciences, 89(16):7452–7456, 1992.

[16] Yu-Sheng Cong et al. The human telomerase catalytic subunit htert: organization of the gene and characterization of the promoter. Human molecular genetics, 8(1):137–142, 1999.

[17] Wyeth W Wasserman et al. Applied bioinformatics for the identification of regulatory elements. Nature Reviews Genetics, 5(4):276–287, 2004.

[18] Matthew E Ritchie et al. limma powers differential expression analyses for RNA-sequencing and microarray studies. Nucleic Acids Research, 43(7):e47, 2015. doi: 10.1093/nar/gkv007.

[19] Meena Choi et al. Msstats: an r package for statistical analysis of quantitative mass spectrometry-based proteomic experiments. Bioinformatics, 30(17):2524–2526, 2014. doi: 10.1093/bioinformatics/btu305.

[20] Xiaofeng Zhang et al. Proteome-wide identification of ubiquitin interactions using ubia-ms. Nature Protocols, 13(3):530–550, 2018. doi: 10.1038/nprot.2017.147.

[21] Samuel A Lambert et al. The human transcription factors. Cell, 172(4):650–665, 2018. doi: 10.1016/j.cell.2018.01.029.

[22] Yulia A Medvedeva et al. EpiFactors: a comprehensive database of human epigenetic factors and complexes. Database, 2015:bav067, 2015. doi: 10.1093/database/bav067.

