## Supplemental Figures for "GLproxScape reconstructs spatial chromatin occupancy landscapes from tiled genomic locus proteomics"

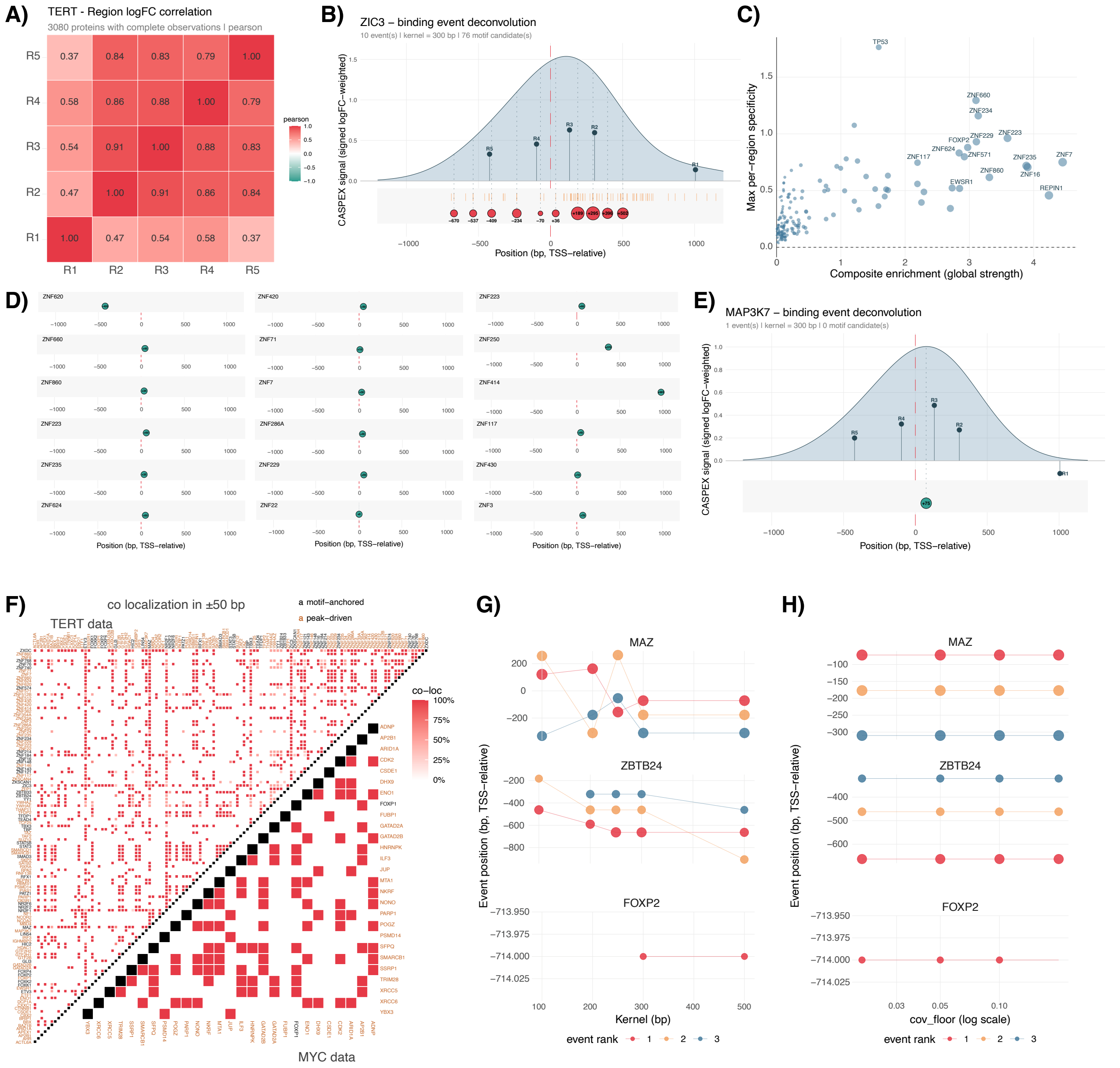


**Supplemental Figure 1: Supplementary analyses of the TERT genomic region.** A) Pearson correlation of protein enrichments of 5 regions targeted with gRNAs in dataset. B) Motif-anchored binding predictions for ZIC3 in TERT genomic region. C) Global strength and per-region specificity of enriched TFs. D) Motif-free peak prediction of ZNF factors in the dataset. E) Motif-free peak prediction of MAP3K7. F) Co-localization of predicted binding events in TERT and MYC genomic region datasets. G-H) Event position shifts of MAZ, ZBTB24, and FOXP2 with different kernel (G) and coverage floor (H) settings in GLproxScape.

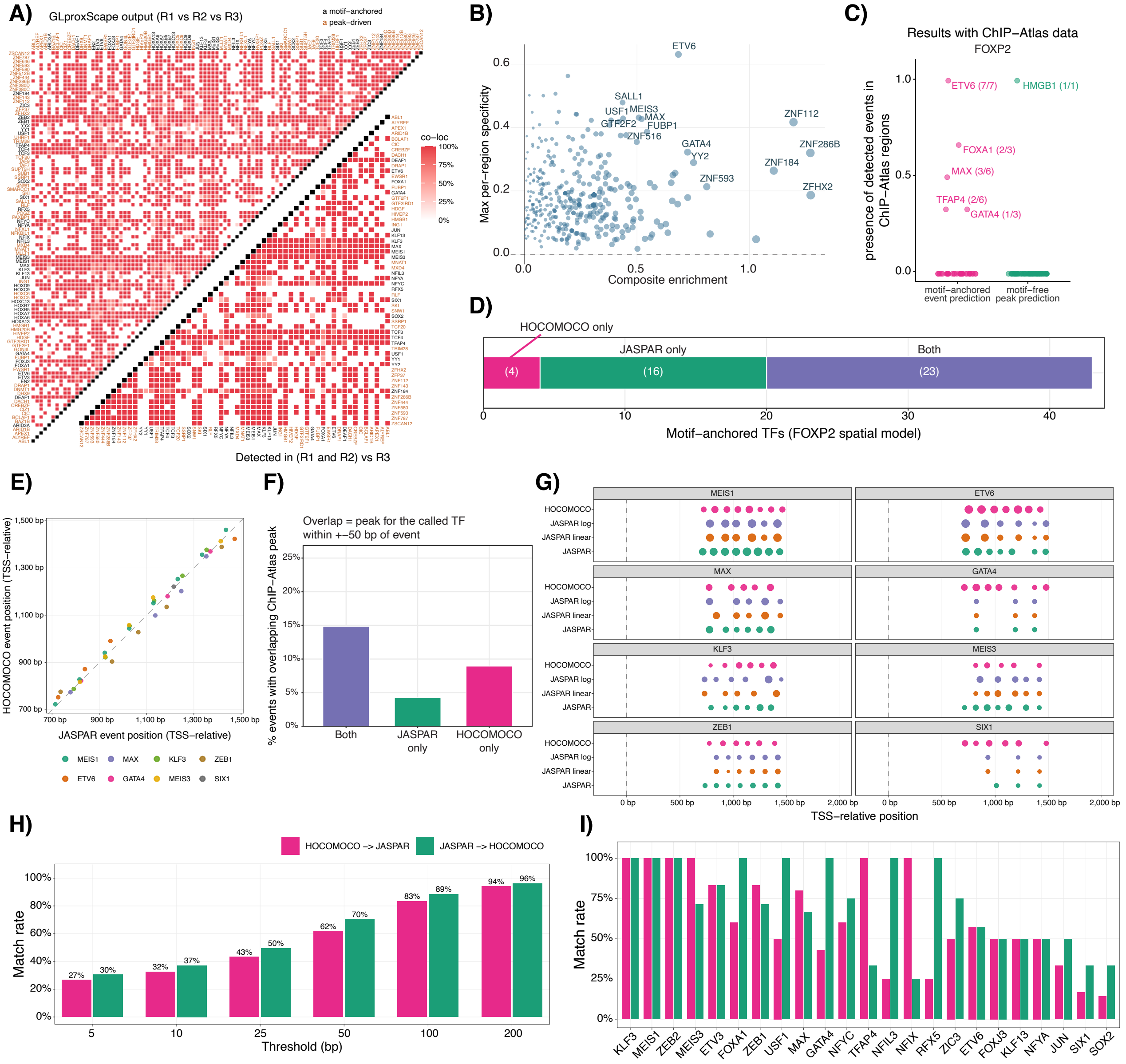


**Supplemental Figure 2: Supplementary analyses of the FOXP2 genomic region.** A) TF-pair co-localization of predicted binding events at the FOXP2 region. Because R1 and R2 overlap by ∼15 bp (effectively the same region sampled twice), the standard co-localization map is inflated by single-region-only signals at that position (top-left panel). Restricting to proteins detected at both R1 and R2 before scoring co-occurrence yields fewer but more biologically interpretable pairs (bottom-right panel). B) Global strength and per-region specificity of enriched TFs. C) The coverage ratio of predicted events in ChIP-Atlas data (matched within ± 25 bp) in FOXP2 dataset. D) Counts of motif-anchored binding predictions with JASPAR and HOCOMOCO databases in GLproxScape. E) Comparison of event positions in GLproxScape analysis with JASPAR and HOCOMOCO. Top eight TFs by combined NNLS amplitude ($\beta$), restricted to TFs called motif-anchored by both databases, were plotted. F) Percent events overlapping with ChIP-Atlas peaks with JASPAR and HOCOMOCO analyses. G) Event positions of the same eight TFs as panel E with different JASPAR weight settings and HOCOMOCO in GLproxScape analysis. H) Match rate vs nearest-neighbour threshold for motif-anchored events. For each event called by one database, the nearest-neighbour distance to a same-TF call in the other database was computed without an a priori matching gate; cumulative match rate is shown at thresholds of ±5, 10, 25, 50, 100, and 200 bp, separately for the JASPAR → HOCOMOCO and HOCOMOCO → JASPAR directions. I) Per-TF match rate at ±50 bp for the 23 TFs called motif-anchored by both databases.

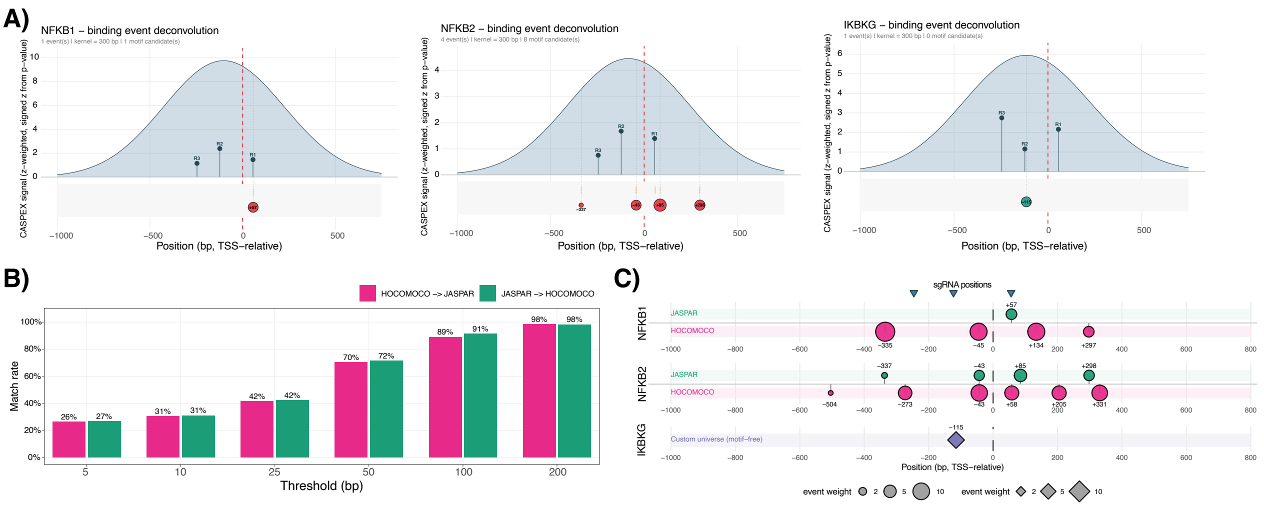

**Supplemental Figure 3: GLproxScape analysis of Ripk3 genomic region.** A) GLproxScape predictions for NFKB1 and NFKB2 (motif-anchored via JASPAR) and IKBKG (motif-free, recovered from a custom run with modified TF universe). B) Cross-database concordance between JASPAR and HOCOMOCO mouse motif-anchored event calls at the Ripk3 promoter: nearest-neighbour match rate at thresholds of ±5, 10, 25, 50, 100 and 200 bp, reported separately for the JASPAR → HOCOMOCO and HOCOMOCO → JASPAR directions. C) Side-by-side comparison of motif-anchored binding events called by JASPAR and HOCOMOCO mouse for NFKB1 and NFKB2, plus the motif-free IKBKG peak from the custom-universe run. Bubble size encodes event weight.

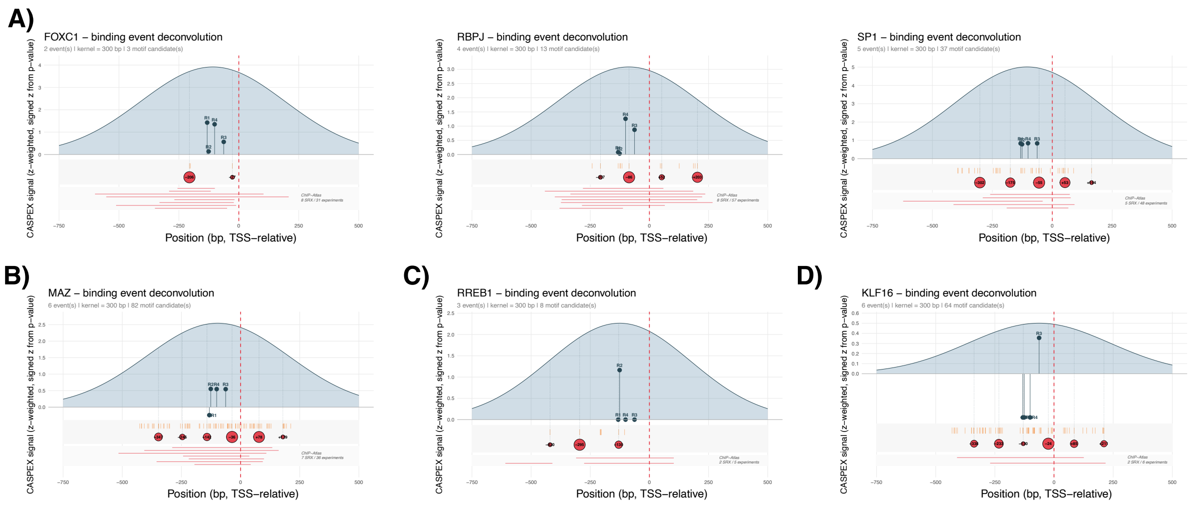


**Supplemental Figure 4: GLproxScape analysis of FOXQ1 genomic region.** A) Motif-anchored binding-event predictions for FOXC1, RBPJ, and SP1 at the FOXQ1 promoter. B) Motif-anchored prediction for MAZ, showing multi-region recovery across the FOXQ1 promoter. C) RREB1 prediction, recovered despite strongly imbalanced per-region signal. D) KLF16 prediction, recovered in a more extreme imbalance case where three sgRNAs showed explicitly negative enrichment and only one sgRNA contributed with weak logFC positive signals.
